# An interactive 3D atlas of gene expression in the mouse brain

**DOI:** 10.64898/2026.01.20.700446

**Authors:** Harry Carey, Sébastien Piluso, Ingvild Bjerke, Xiaoyun Gui, Sharon Yates, Sophia Pieschnik, Signý Benediktsdóttir, Archana Golla, Maja Puchades, Timo Dickscheid, Trygve Leergaard, Simon McMullan, Henry Markram, Daniel Keller, Jan G. Bjaalie

**Author notes:** Correspondence: Jan G. Bjaalie Harry Carey. Co-first authorship (Harry Carey, Sébastien Piluso).

## Abstract

The Allen Mouse Brain Atlas provides gene expression data for over 20,000 genes and has been extensively used in neuroscience. However, it’s constrained by a resolution of 200 µm^3^. We improved this to 25 µm^3^ for 4,083 genes by re-registering 401,660 sections to atlas space. From this, we produced an atlas with regions defined by shared molecular signatures and built an interactive online platform allowing exploration of gene expression patterns.

## Main

Gene expression atlases have substantially advanced our understanding of the organization of the mouse brain^1–3^. Of particular impact has been the Allen Mouse Brain Atlas (AMBA), offering brain-wide expression data for over 20,000 genes across the C57BL/6 (standard laboratory mouse) whole brain^1–4^. This resource is based on classical *in situ* hybridization (ISH) experiments, where the expression of each gene is measured in histological sections. Sections are registered to the three-dimensional (3D) Common Coordinate Framework version 3 (CCFv3) mouse brain atlas^5^, and reconstructed into 3D maps of gene expression. The AMBA has supported discoveries across neurodegeneration, neuroplasticity, and genetic disorders^6^. However, the AMBA is constrained by relatively coarse spatial resolution^7^ (200 µm^3^) and contains registration inaccuracies^8^ which blur anatomical detail.

Recent advances in registration methods now allow for more accurate alignment of histological sections to 3D reference atlases than what was possible when the AMBA was first developed^8–11^. Using the DeepSlice toolkit^8^, along with the Advanced Normalisation Tools (ANTs) nonlinear alignment tool^9^, we re-registered the coronal portion of the AMBA ISH data (4,083 genes) to an extended version of the CCFv3, called the CCFv3 Blue Brain Project (CCFv3_BBP_)^12^. We then combined these registrations and segmentations to generate 3D gene expression volumes at 25 µm^3^, a substantial increase over the 200 µm^3^ of the AMBA. To support exploration and reuse, the data has been shared through the EBRAINS Knowledge Graph (RRID:SCR_017612) and volumes have been integrated into the siibra explorer atlas viewer^13^.

The main advance of this work is that it provides far more detailed spatial information than was previously possible^9^. By reconstructing thousands of datasets at high resolution in a common coordinate framework, the resource enables more refined and intuitive exploration of gene expression patterns across the whole mouse brain. It also enables the automated identification of areas based on shared expression signatures.

To create this resource, we began by downloading all coronal ISH datasets from the AMBA which were from postnatal day 56 C57BL/6 mice (totalling 401,660 sections from 1,078 brains). We used DeepSlice (RRID:SCR_023854) to place each section into the CCFv3_BBP_. DeepSlice has previously been shown to achieve expert level section image to atlas registration accuracy^8^. For each brain, a subset of the alignments were reviewed and corrected with QuickNII (RRID:SCR_016854)^10^, and any changes to angle or position were propagated to all sections from that brain. Nonlinear tissue distortions were corrected using the Advanced Normalisation Tools (ANTs) toolbox, which registered each section to the corresponding Nissl template slice from the CCFv3 _BBP_ ^12^. This procedure was repeated for all sections (Figure 1a).

**Figure 1.**
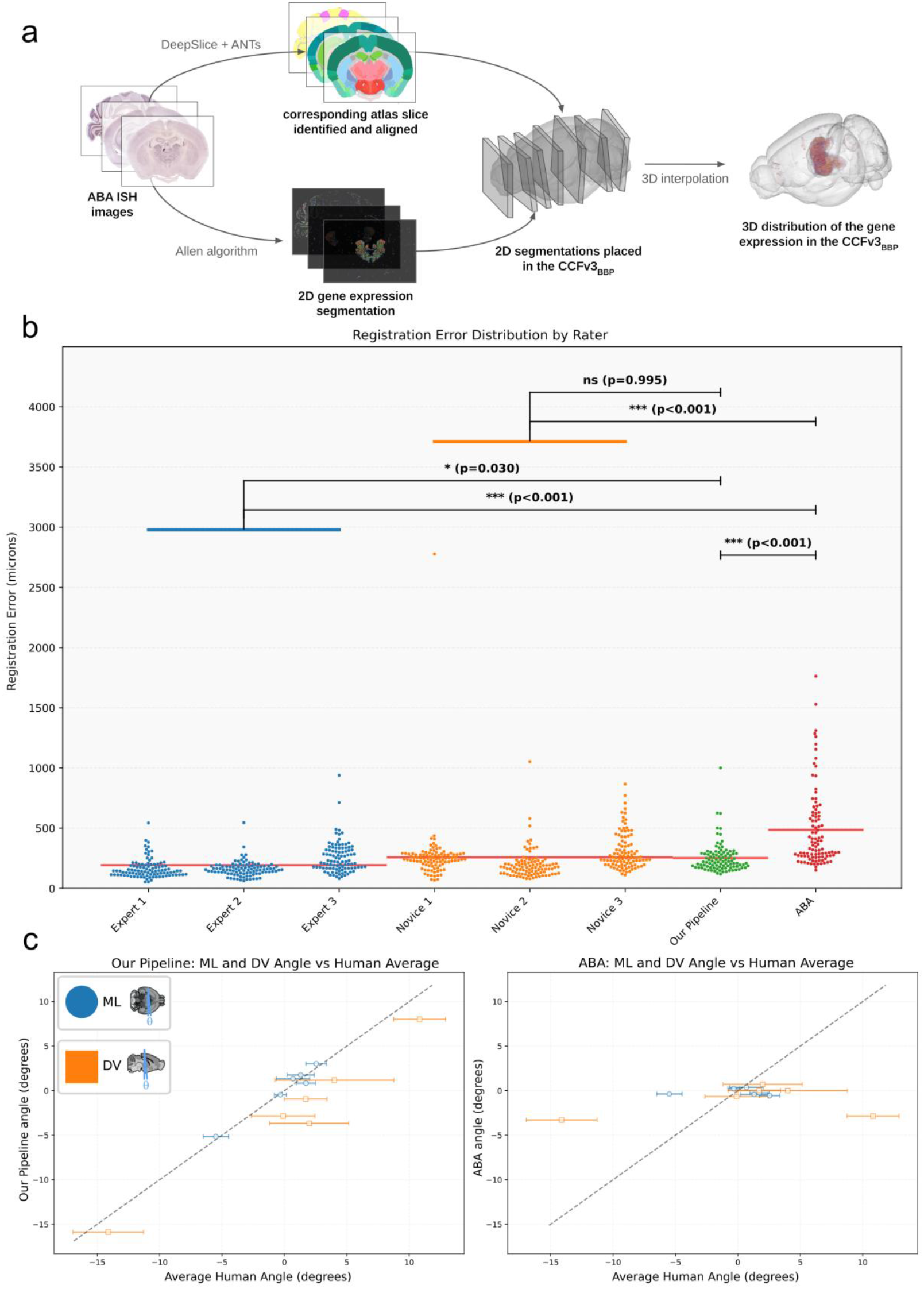
a, Pipeline overview. Coronal ISH sections from the Allen Brain Atlas are registered to the CCFv3bbp reference atlas using DeepSlice. Non-linear distortions are corrected using Advanced Normalisation Tool (ANTs). Empty space between sections is then interpolated creating high resolution 3D gene expression maps. b, Quantification of registration error. The plot shows the distribution of alignment errors for novices (n=3), experts (n=3), our automated pipeline, and the original AMBA alignments. Our pipeline significantly outperforms the AMBA (p<0.001) and performs at a level comparable to novices (p=0.995). Red lines indicate the mean error for each group. Each data point represents one section c, Comparison of predicted sectioning angles. Scatter plots show the correlation between the average human-annotated angles (x-axis) and the angles predicted by our pipeline (left) and the original AMBA (right) for mediolateral (ML, blue) and dorsoventral (DV, orange) axes. Our pipeline’s predictions show a strong correlation (r2=0.823) with human annotations, unlike the AMBA’s (r2=0.03).

To validate our registration pipeline, we compared its accuracy with the original AMBA registrations and with manual alignments (both from experts and novices) generated using QuickNII and VisuAlign (RRID:SCR_017978)^11^ (figure 1b). For the manual alignments, each rater placed sections from nine AMBA brains into the CCFv3_BBP_ using QuickNII, and performed non-linear corrections for distortions using VisuAlign. We then measured how far each rater’s registrations within the CCFv3_BBP_ deviated from the group average, providing an estimate of alignment error in microns^8^. We repeated this process, measuring the deviations of our alignments, and the AMBA’s, from this group average.

Experts were most accurate (mean error: 193.6 µm), with novices being significantly less accurate (mean error: 257.9 µm; Tukey *P*<0.001). Our pipeline matched novice performance (mean error 252.8 µm; Tukey *P*=0.995), while remaining significantly less accurate than experts (Tukey *P*<0.05). Our pipeline outperformed the original AMBA registrations (mean

AMBA error 485.9 µm; Tukey *P*<0.001), as did novices (Tukey *P*<0.001) and experts (Tukey P<0.001). A major limitation of the AMBA registrations was poor estimation of cutting angles, which showed little correlation with those determined by neuroanatomists (r2 = 0.03, figure 1c). By comparison, the angles specified with our pipeline were highly correlated with the average neuroanatomist-determined angle (r2 = 0.823, Figure 1c).

Since our pipeline produced significantly more accurate alignments than those in the AMBA, we were able to construct gene expression maps at much higher resolution than the original 200 µm^3^ volumes provided by the Allen Institute. Using these improved alignments, we generated 25 µm^3^ 3D volumes for 4,083 genes. Gaps between sections were interpolated, and experiments targeting the same gene were combined into a single averaged volume. This increase in resolution greatly improved visualization of gene expression in fine subregions (figure 2A). Genes with many available experiments such as Cap1 and Cacna1g (top two panels of figure 2A) yielded the most detailed reconstructions. However, even genes with a low number of samples can be reconstructed into high-quality 3D maps using our approach (bottom two panels of figure 2A). Since many datasets also contained Nissl stained sections, these were also registered and combined into a population average Nissl volume, this volume is reported on in more depth in Piluso et al.^12^.

**Figure 2.**
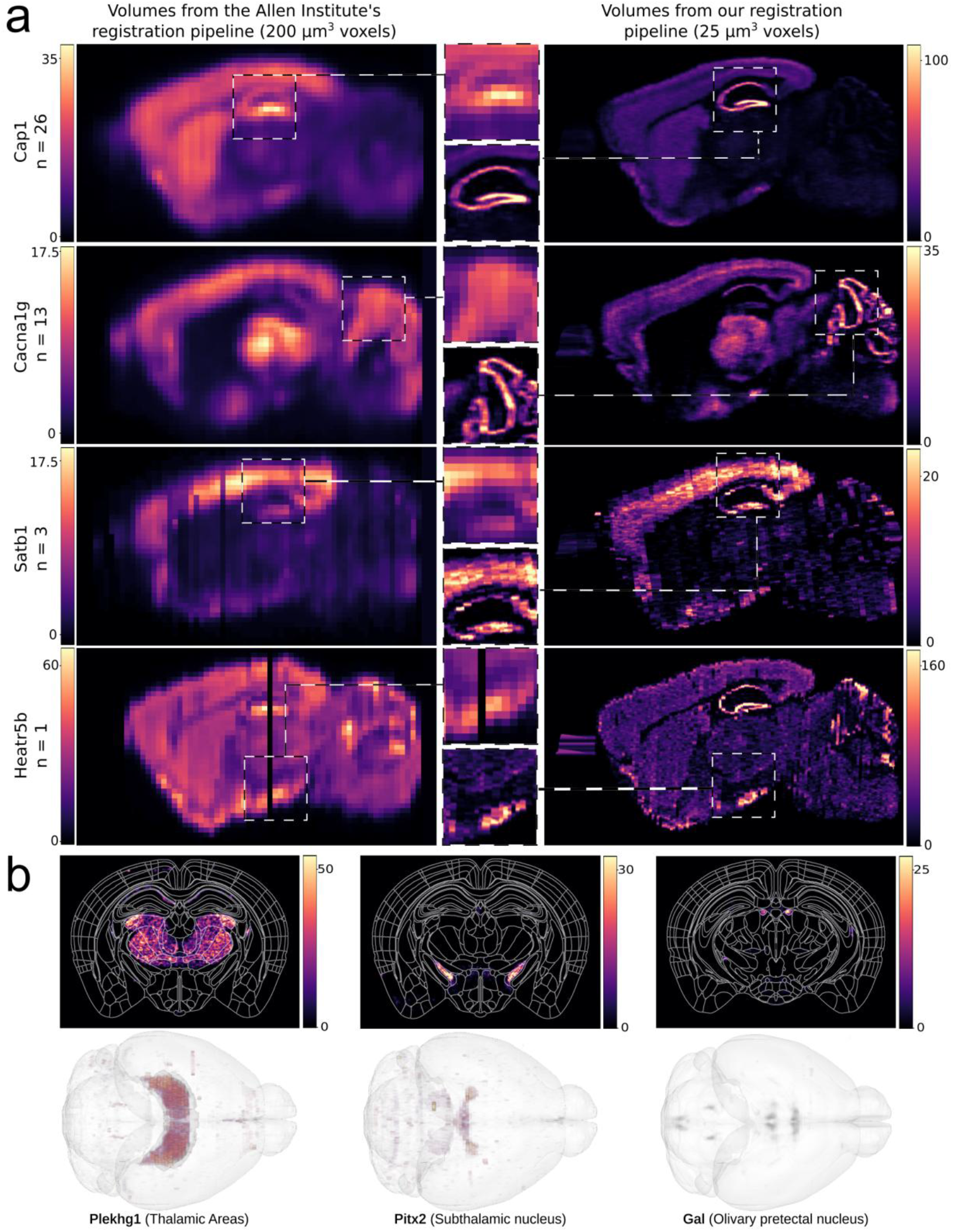
a, Comparison of gene expression volumes from the original Allen Brain Atlas (left, 200 µm^3^ voxels) and our pipeline (right, 25 µm^3^ voxels). Our high-resolution maps reveal finer anatomical details for genes with varying numbers of experimental samples (n). From top to bottom: Cap1 (n=26), Cacna1g (n=13), Satb1 (n=3), and Heatr5b (n=1). The values on the colour bar represent the range of shown pixel values. We set the max intensity of each plot at 70% of the max value in the volume so as to better display the variance present in each gene. The Allen institute has a smaller range of pixel values since it is lower resolution, greater down sampling has a smoothing effect which removes outlier values. b, Examples from the spatial search tool. The top-ranked gene is shown for queries targeting thalamic areas (Plekhg1), the subthalamic nucleus (Pitx2), and the olivary pretectal nucleus (Gal). The 3D view (left) shows the expression of Plekhg1 within the thalamus (red) in the context of the whole brain. The coronal views show the precise expression patterns of Pitx2 and Gal within their respective target regions (outlined).

To make the resource more accessible, we built an online portal, available at https://neural-systems-at-uio.github.io/spatial_brain_maps/, where users can search for genes of interest and explore their expression patterns. The portal also lets users select atlas regions and rank genes by specificity of gene expression in the chosen brain region, coverage, intensity, or a weighted mix of these metrics (figure 2b, supplementary figure 1). For interactive 3D viewing, we integrated all volumes into the siibra-explorer^13^ on the EBRAINS platform, allowing each gene to be visualised alongside an atlas.

For more than a century, brain atlases have been shaped by the methods available for defining neuroanatomical boundaries. Early atlases relied on features visible under the microscope through histochemical staining, later supplemented by functional criteria. Today, high-resolution spatially registered datasets offer a way to automatically discover brain regions from patterns of gene expression. Earlier efforts have used clustering of AMBA data to create regions based on differential gene-expression^14–17^, but these attempts were limited by the original 200 µm^3^ AMBA resolution. Our volumes overcome this constraint, with 25 µm^3^ resolution allowing clustering at a resolution comparable to contemporary 3D atlases^5^. Using our gene-expression volumes we first mirrored one hemisphere onto the other to create a symmetrical dataset. The dimensionality of the data was reduced with Principal Component Analysis (PCA) and K-means clustering was applied, grouping voxels with similar expression profiles. The resulting clusters were then projected back into 3D to generate a fully data-driven atlas.

To select the number of clusters, we tried various numbers of clusters on a subset of voxels and identified an “elbow point” where adding more clusters produced only small gains in performance (supplementary figure 2). This led us to choose 55 clusters for the final atlas. We refer to the final volume as the clustered area (“CArea”) atlas and treat each cluster as a region. Many regions in the CArea atlas recapitulate established neuroanatomical boundaries, such as the cortical layers (figure 3b, slices 186, 225, 286, and 379), and the border between the caudate-putamen and accumbens (figure 3b, slice 379). Some of these regions have been difficult to define in traditional murine brain atlases^18^, highlighting the value of a method which is able to identify these boundaries more objectively.

**Figure 3.**
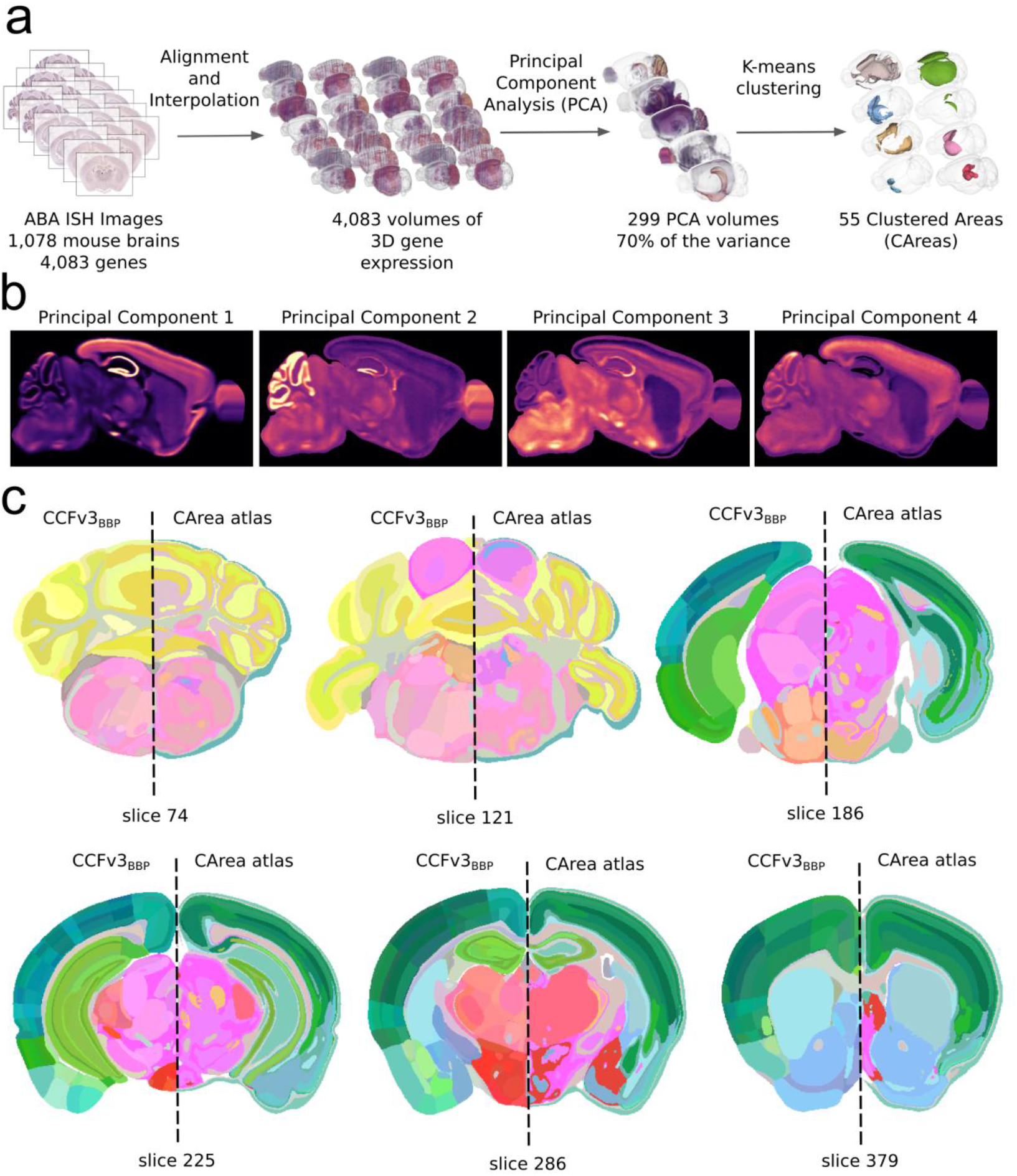
a, Sagittal views of the top 4 principal components after running a voxel-wise iterative PCA on all 4,083 gene volumes. The colour scale represents the principal component (PC) value with bright yellow indicating high positive values and dark purple indicating low or negative values. These components capture the dominant patterns of co-expression across the mouse brain b, Side by side comparison of the regions from an established anatomical atlas (CCFv3_BBP_, left of dashed line) and those from our data-driven Clustered Area atlas (CArea atlas, right of dashed line) on representative coronal slices. The CCFv3_BBP_ colours were modified such that each region was uniquely coloured. Each CArea was assigned the colour of the CCFv3_BBP_ region it most overlapped with, if two CAreas were assigned the same colour, one was adjusted such that each CArea was uniquely coloured. Slice numbers refer to the coronal plane of the atlas.

While spatial transcriptomic methods now allow the simultaneous imaging of thousands of genes, they are not yet scalable to the large cohorts needed for population-level atlasing^19–21^. Our resource integrates data from 1,078 animals, making the CArea atlas regions broadly representative rather than driven by a few mice. Since the atlas is in 3D, it is compatible with existing volumetric atlas analysis software^22–24^. Given that the number of clusters output from our pipeline is adjustable, it is possible for users to create use-case specific atlases if a particular number of parcellations is required. We consider the CArea atlas to be complementary to existing human-delineated atlases such as the CCFv3.

In summary, we present high-resolution 3D gene expression volumes for the mouse brain. The volumes enable a population-scale, data-driven anatomical atlas that captures shared molecular signatures while faithfully recapitulating many known anatomical structures.

Beyond the core dataset, we provide an intuitive search and visualization interface, ensuring accessibility. The full analysis pipeline is reusable and can be adapted to a wide range of future applications, bridging high-resolution molecular mapping with practical tools for exploration and investigation.

## Data sharing statement

- Our online search portal can be accessed at https://neural-systems-at-uio.github.io/spatial_brain_maps/.
- All code for replicating our analysis is available from https://github.com/Neural-Systems-at-UIO/interactive_gene_expression.
- All registrations, gene expression volumes, and clustered atlases are openly accessible via the EBRAINS Knowledge Graph from https://search.kg.ebrains.eu/instances/7f8ef0e2-121a-4892-8a5e-1c7a8b693503.
- The CArea atlas will be released via the brainglobe atlas API (RRID:SCR_023848)^24^.

## Acknowledgements

We thank Heidi Kleven for her invaluable insights into atlasing terminology and methods, Lydia Ng for early discussions that helped shape the project’s direction, and Michael Hawrylycz for valuable comments on the manuscript. We are also grateful to the Allen Institute for Brain Science for sharing their data in a curated and usable form. Finally, we acknowledge Sergio Rivas-Gomez from the Blue Brain Project for his high-performance computing support. This study was supported by the European Union’s Research and Innovation Program Horizon Europe under Grant Agreement no. 101147319 (EBRAINS 2.0), The Research Council of Norway under Grant Agreement no. 333157 (Norwegian INCF Node), the Blue Brain Project, a research centre of the École polytechnique fédérale de Lausanne (EPFL), and the Swiss government’s ETH Board of the Swiss Federal Institutes of Technology.

## Online Methods

### Data acquisition

All ISH and corresponding segmentation data were downloaded via the Allen Institute’s API (RRID:SCR_005984) and filtered to include only coronally cut mouse brains which passed quality control and were from postnatal day 56, C57BL/6 mice. We used the CCFv3_BBP_ atlas^9^ as the anatomical reference. This atlas is an extended and refined version of the Allen Institute’s CCFv3 atlas, incorporating several important improvements. It includes a non-truncated main olfactory bulb, cerebellum, and medulla, thereby providing coverage of the entire mouse brain. It also offers more detailed anatomical annotations, including of the laminar organization within the cerebellum. In addition, the CCFv3_BBP_ provides a 3D Nissl-stained reference volume that is more accurately co-registered with the atlas labels than the Nissl volume provided with the CCFv3.

### Data registration

To register the data, we first ran each whole brain dataset through DeepSlice (version 1.2.4), using the angle integration options and providing the section thickness reported by the Allen so as to improve accuracy. From each dataset, 5 sections covering the length of the brain were selected for quality control (most brains contained between 100 and 500 sections) and reviewed to correct errors in angle estimation or anteroposterior positioning. The cross-section from the CCFv3_BBP_ Nissl reference template corresponding to each recalibrated ISH section was used for non-linear refinements. Using this Nissl section as reference image and the corresponding ISH histological tissue as the moving image, we applied the ANTs registration package and calculated both affine and non-linear transformations.

### Interpolation

Segmentation images were first downsampled via average pooling to match the resolution of the CCFv3_BBP_ (25 µm^3^). We then placed the segmentation data into CCFv3_BBP space_ using the position as determined by our registration files. To correct for empty space between the sections we used a K nearest neighbour (KNN) interpolation method, where each voxel was assigned the average value of the five nearest segmentation pixels. To ensure that all voxels in each volume were equivalent and comparable we applied KNN approach to all voxels not just those for which segmentation data was missing, avoiding a scenario where some voxels are an average of 5 and others just single datapoints. This process produced smooth 3D volumes of gene expression at 25 µm^3^ resolution.

### Validation

To quantify alignment error for each section, raters positioned sections in atlas space using QuickNII and VisuAlign. The registration generated by each rater was then compared to the mean position derived from all other raters. To obtain an error measure that reflected the full extent of each section rather than only its edges, a 2D grid was projected into the 3D atlas space. Every grid point was transformed according to the rater’s placement, and an identical grid was generated using the mean placement of the remaining raters. The Euclidean distance between corresponding points in these two grids (supplementary figure 3), expressed in microns, was used as the section-wise error metric (Figure 1b).

### Clustering

In order to make clustering across all 4,083 genes computationally feasible, we first ran a voxel-wise iterative Principal Component Analysis (PCA) on the gene volumes in order to reduce the dimensionality down from 4,083 genes per voxel to 299 principal components, retaining 70% of the original variance. K-means clustering was then applied to this reduced data, clustering together voxels with similar molecular signatures. We then projected these clusters back into 3D producing atlas like volumes identifying regions of the brain with similar patterns of gene expression. The K-means clustering algorithm allowed us to choose the number of clusters, we therefore tested cluster numbers from 10 to 100 (incrementing by 5) on a 1,000,000 voxel subset (supplementary figure 2). By analysing the change in the silhouette score, we were able to manually identify an elbow point in the graph at 55 clusters. For this reason, we chose the 55 regions for our “CArea” (Clustered-Area) atlas (each cluster functions as a region of the CArea atlas).

---

**Supplementary Figure 1.**
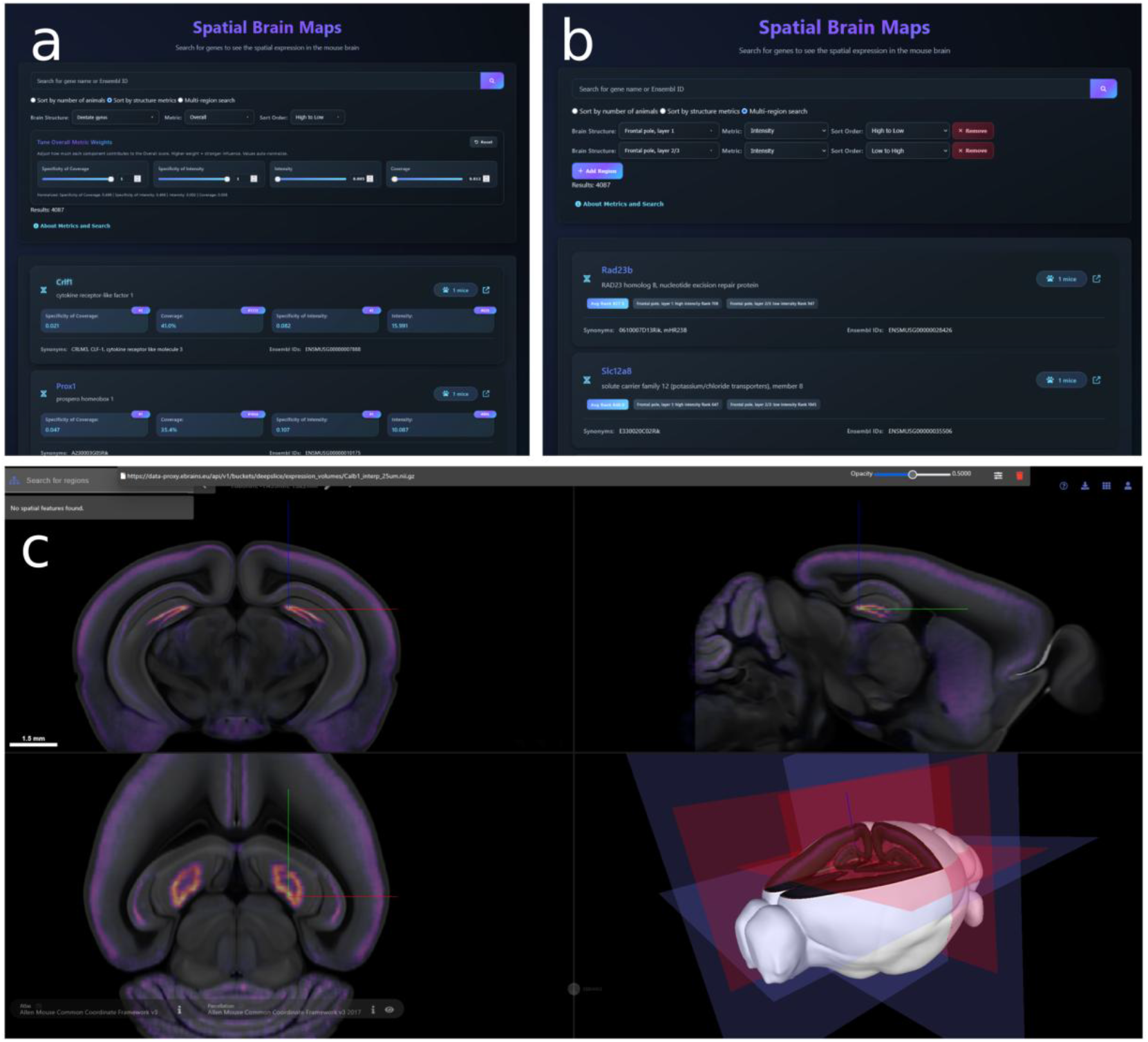
a, The region based search interface for our dataset. The input allows you to choose a region and then weight the ranking according to an array of metrics. b, with a more advanced multi region search functionality users can find genes which are highly expressed in one region but lowly expressed in another. Users can combine as many regions as they wish into their query. c, All volumes are viewable in 3D in the browser via the Siibra atlas explorer.

**Supplementary Figure 2.**
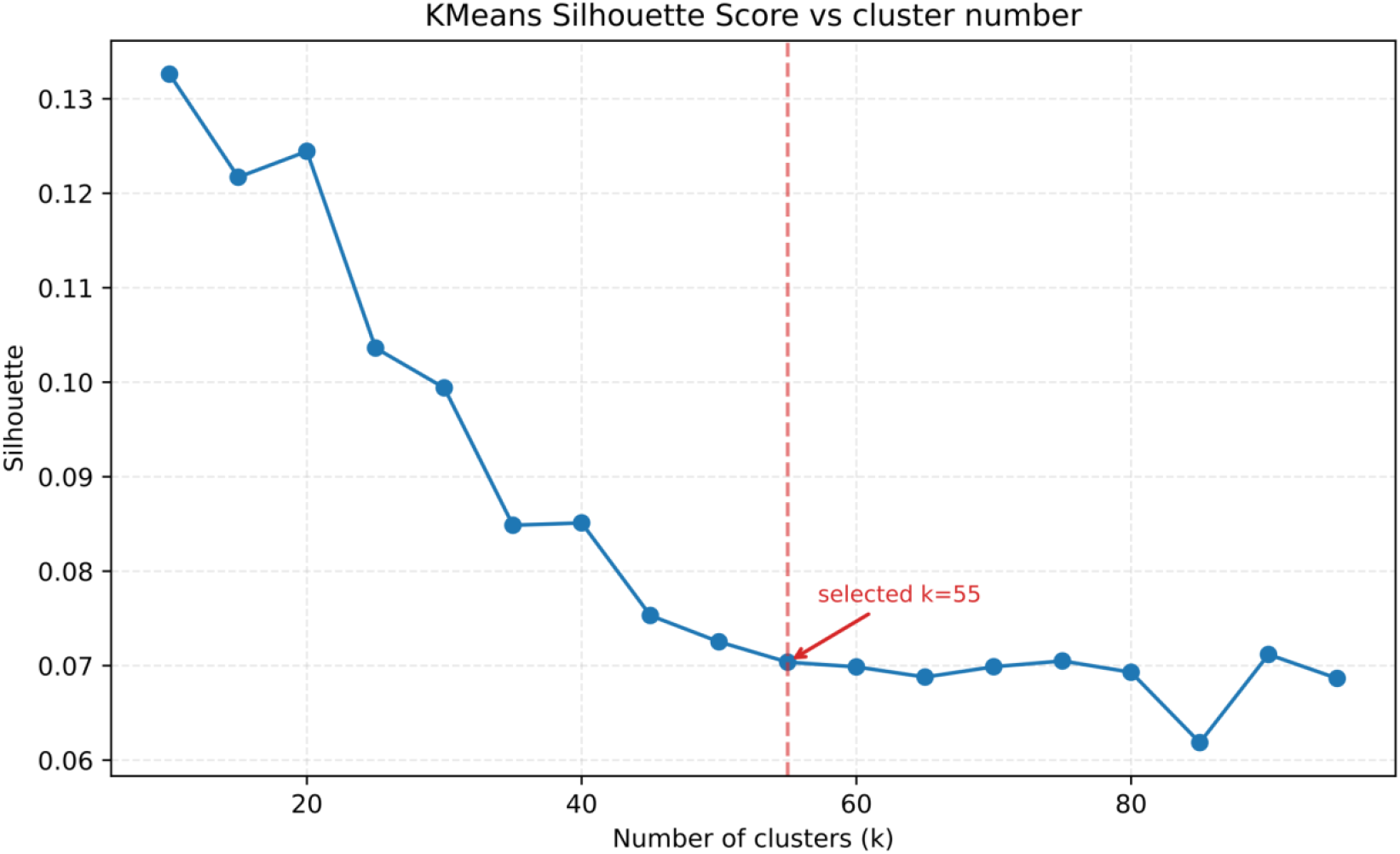
Running the k-means algorithm on a subset of the PCA voxels (1,000,000 voxels) helped us identify an elbow point at 55 regions. Due to this we chose 55 as the number of regions in the CArea atlas.

**Supplementary Figure 3.**
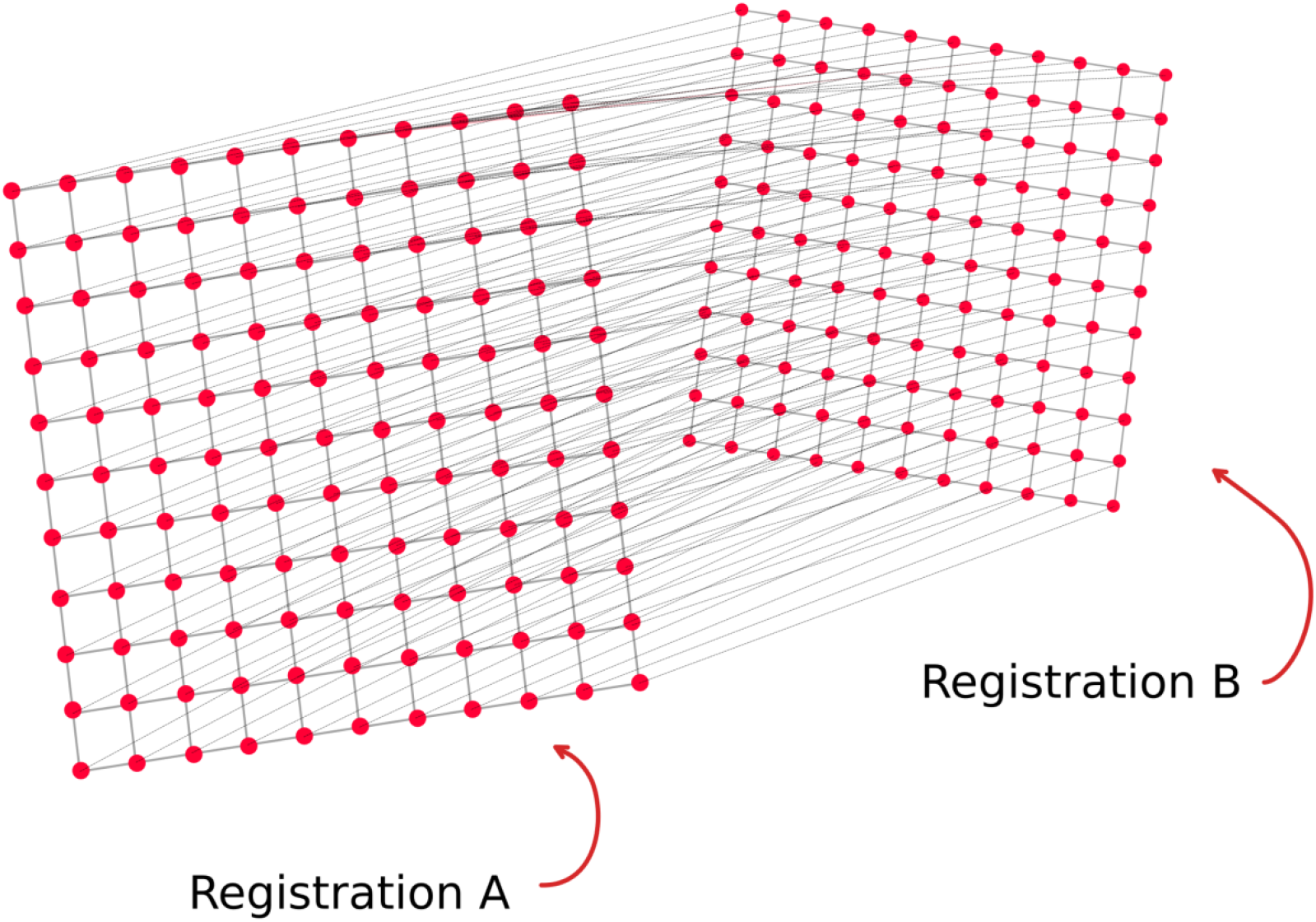
To measure the distance between two registrations we project a grid into atlas space, where the corners of the grid align with the corners of the registration. We then measure the pairwise distance between corresponding points in each grid.

